# Grid cells are not a parsimonious source of the hexadirectional signal

**DOI:** 10.64898/2026.03.04.709667

**Authors:** Noam Almog, Thanh Pierre Doan, Tobias Navarro Schröder

## Abstract

Hexadirectional modulation is widely treated as a proxy for population-level grid-cell activity in humans. Conventional analyses test the sixth harmonic of the mean directional response, whereas we show grid-cell firing predicts a dominant sixfold modulation of cell-level variance, rather than population-level mean. Using analytical derivations, simulations and recordings from 23,453 rodent medial entorhinal neurons, we show that grid cell tuning alone cannot account for the measured mean signal. Analysis of 1,349 conjunctive grid-head-direction cells revealed insufficient grid-axis alignment and directional tuning specificity to account for population-level hexadirectionality. A hypothetical additional nonlinearity mechanism near the cell level can produce a a small sixfold population-level mean modulation, ≤ 2%, under favorable circumstances. Moreover, the current methodology does not adequately differentiate genuinely symmetric activity and stable directional peaks, nor contextualize a harmonic measurement in a solid spectral framework. These findings show that mean hexadirectional modulation neither follows directly from grid-cell firing nor uniquely identifies grid-cell activity. We provide an improved framework for testing a link between the two in future studies.

## Introduction

A central goal in cognitive neuroscience is to understand how the emergence of the mind and cognitive faculties such as memory relate to the activity of cells in the brain. One of the most successful research fields pursuing the goal of linking activity of single neurons to clearly defined cognitive variables has been that of spatial navigation. Head direction cells, mainly found throughout brain regions of the Papez circuit, are tuned to the azimuth movement direction in the horizontal plane^1,2^. While place cells in the hippocampus and grid cells in the medial entorhinal cortex (MEC) are tuned to the spatial location of an animal^3–5^. These striking findings were first made in freely moving rodents. Linking cognitive variables of spatial location to the activity of single neurons at the top of the cortical hierarchy far removed from primary sensory areas was recognized as a breakthrough worthy of a Nobel Prize^6^. Testing the relevance of these findings for the human brain and higher cognition was hampered by the necessary invasive recording procedures.The discovery of an fMRI-based signature of putative grid cell population activity (termed hexadirectional signal or grid cell-like representations) during free navigation in virtual reality experiments^7^ claimed that one could investigate the specific role of grid cells in higher order association cortices non-invasively in humans.

While this method has detected hexadirectional most often in the entorhinal cortex, significant sixfold effects have also been reported repeatedly in vmPFC^8–12^ and in anterior piriform cortex during olfactory navigation^8^, while broader whole-brain analyses have additionally implicated posterior cingulate/precuneus, retrosplenial, posterior parietal, orbitofrontal and temporoparietal regions^7,9,11^. Behaviorally, the hexadirectional signature has been reported during actual and imagined navigation^7,13^, conceptual, olfactory, semantic and social spaces^8,9,11,14^, visual exploration and memory^10,15,16^, and experimentally induced changes in perceived self-location^17^. Yet the signal is not consistently detected in EC: significant modulation has been reported during observation but not self-navigation within the same participants^18^, while a recent study us failed to detect the expected group-level EC effect^19,20^.

Crucially the hexadirectional signal is a function of trajectory direction. Whether the single-unit 2D grid cell firing pattern produces the hexadirectional mean population signal remains unknown. Three main hypotheses for a potential mechanisms have been proposed^21^. H1: *Basic geometry*. The 2D grid cell pattern itself leads to activity differences across movement directions. H2: *Conjunctive cell alignment*. Conjunctive grid-x-head-direction cells have a preferred directional tuning relative to grid pattern axes^7,22^ H3: *Non-linear transformation*. Grid cell firing is nonlinearly transformed prior to the measurement of the population signal.

Using analytic derivations, computational simulations, and recordings from 23,453 MEC cells across 29 recordings^23^, we have provided a first-principles explanation for how a statistically detectable population level hexadirectional pattern can arise directly from the geometry of the single cell grid pattern itself, namely in the activity *variance* at the single-cell level. This highlights a major contradiction in the field, where the primary quantity measured is *mean* directional modulation, which we show requires secondary mechanisms. In this regard, analyzing 1,349 conjunctive grid x head-direction cells we found limited evidence to suggest that the directional tuning of conjunctive grid cells is anchored to the grid axes, the low specificity of this tuning makes it highly unlikely to account for the observed hexadirectional signal, consistent with our finding that the effect is absent in mean empirical activity in our dataset. We also advance theoretical understanding of the nonlinearity hypothesis through both analytic treatment and empirically grounded computational simulations. We find that despite favorable assumptions, a parsimonious population-level signal based on empirical MEC populations produces a relatively small mean hexadirectional signal, around 2%. Lastly we have provided a critical evaluation of the common hexadirectional analysis method and some of the pitfalls it contains. Namely, quantifying a single frequency magnitude outside a spectral analysis framework as well as not differentiating genuinely symmetric activity from a stable asymmetric one. Taken together, these findings bridge previously disconnected theoretical and empirical perspectives, offering a principled mechanistic account of the hexadirectional signal that reevaluates our understanding of grid coding in humans and provides a new foundation for interpreting large-scale directionally modulated dynamics in the entorhinal-hippocampal system.

## Results

To quantify k-fold rotational symmetry in a directional signal we used Fourier (like Bin Khaled et al.^22^) in a manner equivalent to sinusoidal regression in canonical hexadirectional analysis^7^, where angular-frequency amplitude is a proxy for k-fold rotational symmetric modulation (see Methods). **Signal Contribution** is simply normalized amplitude (magnitude), *Â* _*k*_ = *A*_*k*_ / ∑ *A*, as a percentage. **Segment activity** is the summed activity (firing) over a trajectory segment. The **directional signal** (s,θ) is either the mean or variance of segment activity for all segments sampled at direction θ, quantified for single cell, or population-level (summed over cells) activity.

### Symmetry in directional grid cell activity

Here we examine the most basic hypothesis that grid tuning pattern geometry produces a 6-symmetric directional response. For a description of how spatial/directional sampling is implemented see Methods.

#### Mean firing of a spatial firing pattern is not modulated by direction

To detect grid cells, hexadirectional analysis quantifies rotational symmetry in mean directional neural response (see Methods). However a major contradiction exists: principally, the mean firing of a spatially tuned cell, or population, is not directionally modulated (Figure 1B). Consider the spatial activity map as an image (color=activity). If cut into *n* straight parallel slices (trajectory segments), with activity meaned over slices, the mean remains constant regardless of cutting direction, since the slices always partition the same total activity. Figure 1C reproduces this principle via computational simulation (mean=red). Note that unlike in Figure 1B, simulated segments have same length. Consequently, canonical hexadirectional analysis of mean directional response cannot detect grid cell activity (single cell, or population) from a first-principle perspective, without the presence of additional second-order phenomena.

**Figure 1:**
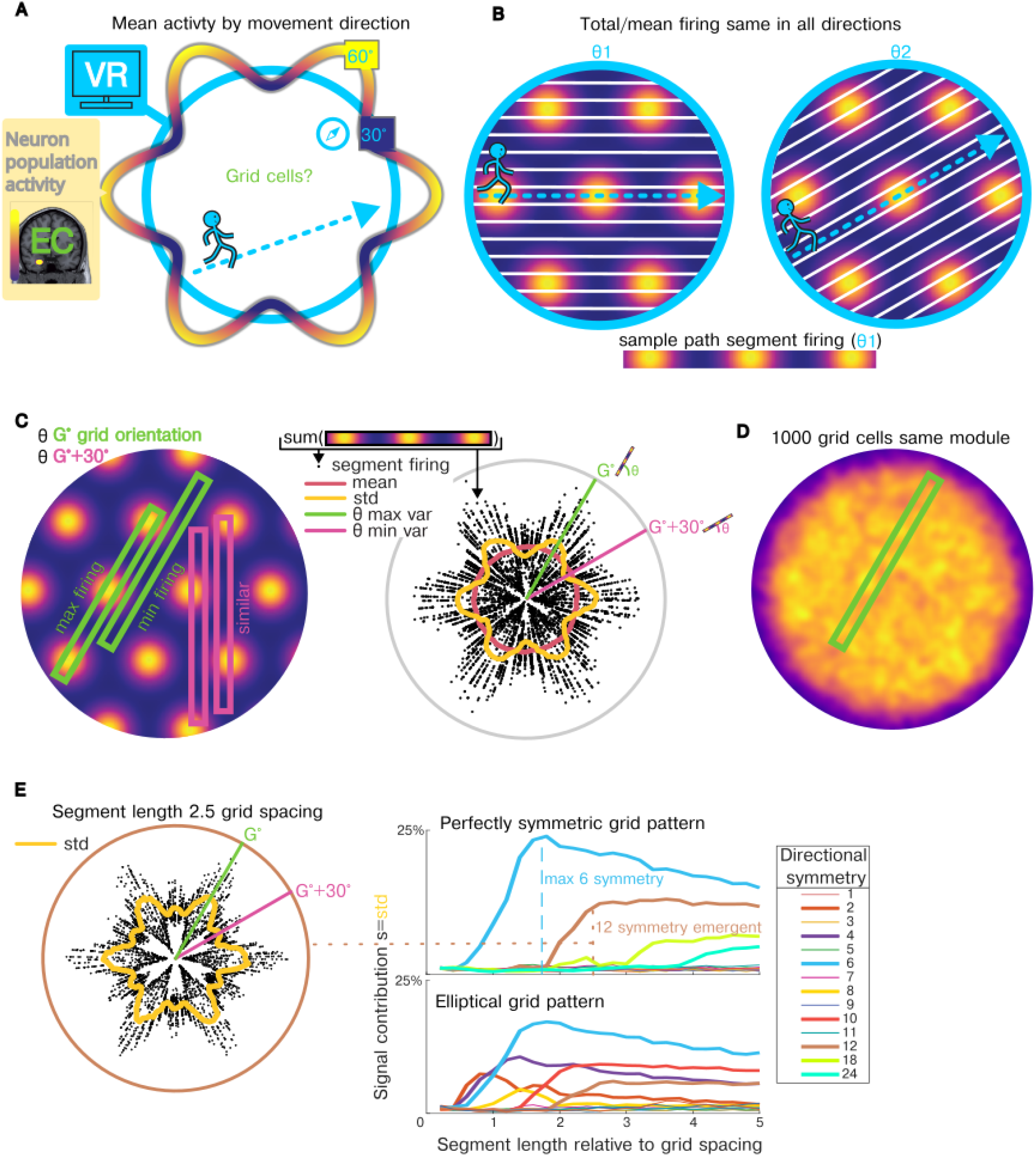
Directional activity produced by grid pattern tuning. (A) Schematic of the hexadirectional signal: 6 symmetric (60°) activity peaks. Reported in human entorhinal cortex recordings (typically population-level, commonly fMRI-BOLD), while participants navigate virtual environments. Hypothesized as the signature signal of a grid cell population. (B) When sampled completely (all locations in a 2D space), *any* spatial tuning pattern produces the same amount of mean/total activity, regardless of sampling direction, because activity summed over the space is constant. Conversely, individual trajectory segment activity like the one here containing 3 fields may only be achievable at specific directions. (C) Unlike mean activity per segment, **variance** is directionally modulated. For an idealized grid pattern, min and max segment activity occurs at grid orientation directions for equal length segments. Right: simulated segment activity over a grid pattern (equal lengths, black points), r-axis=activity. Note how mean is was constant. (D) A spatial tuning map produced by 1000 grid cells with same orientation and grid-espacing and uniform random phase (i.e. same module + small noise). Note how contrast in low/high firing firing is much less than for single cell. (E) Effect of segment sample length on directional variance. Left: for ideal grid pattern geometry, when segment length is 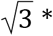 *grid spacing* two fields can be reached at both ‘aligned’ (grid-orientation) and ‘misaligned’ (orientation+30°) directions (right panel, blue dashed-line). This second peak at grid-orthogonal directions produces a harmonic 12-symmetry which accounts for an increasing portion of the signal, peaking at segment length ≈ 2. 5 * *grid spacing*, at which point more fields can be reached at additional directions. As segment length increases, total energy in the signal is distributed across more harmonics. Right bottom: for an elliptical grid pattern, another set of even symmetries also account for a portion of the total energy in the signal. “Signal contribution” is the proportionate Fourier amplitude of the standard-deviation of segment activity by direction as a percentage. Polar sinusoid decomposition is used to quantify rotational k-fold symmetry, equivalent to sinusoidal regression in canonical hexadirectional analysis.

Unlike mean, variance of activity can be modulated by direction depending on the spatial tuning pattern. Using the image analogy again, we see that the activity pattern of a single segment may only be possible in certain directions. In figure 1B we see a segment containing all 3 yellow fields is possible for θ1, not θ2 (also note only this direction contains segments with *no* fields). Consequently, Figure 1C left shows how at grid axial directions we get both highest and lowest activity path segments. Figure 1C right, demonstrates how this produces a hexadirectional variance signal.

In large part the magnitude of this variance modulation is driven by the amount of contrast in the spatial pattern itself (the difference between high and low activity regions). While single cell grid patterns are high in contrast, when grid cells with different phases are recorded together, the contrast in the population-level activity decreases (Figure 1D). Empirically, the cells in a grid module have uniformly distributed phases^4^, implying we should not expect significant hexadirectional variance modulation in a grid cell population signal.

Segment sample length also significantly modulates the hexadirectional signal. While it is intuitive to assume that longer segments produce a stronger signal, this is not the case for a pure grid pattern (Figure 1E). Segment length determines how many fields can be reached in a given direction relative to the grid spacing. Due to the 60° geometry, at a distance of 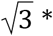 *grid spacing*, two fields can be reached at both the grid-axis directions as well as the orthogonal (‘misaligned’) directions (+30°). This complicates/contradicts the narrative that the ‘misaligned’ directions will have the lowest signal. In Figure 1E we see that at this length we start to see peaks every 30° producing a 12 symmetry. Figure 1E right/top shows that for a pure grid pattern, lengths greater than 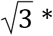 *grid spacing* produce a decreasing amount of 6-symmetry as additional harmonics (multiples of the 6 symmetry) occupy a larger portion of the total energy in the signal. Noisy (empirical) and/or elliptical (Figure 1E right bottom) grid patterns will also decrease the signal. Critically, varying segment sample lengths (as in an openfield trajectory) will also lower the variance signal. Segment lengths shorter than grid-spacing included in analysis only add noise, as the same number of fields (one) can be crossed in all directions.

#### Null distribution of directional symmetries in directionally and spatially tuned population activity

The MEC is also home to other directionally and spatially tuned cells besides grid cells^2,24^ Figure 2A. To understand the null directional signal they produce we simulated random populations of directionally and spatially modulated cells (see Methods) in Figure 2B. Directionally tuned cells produced a null distribution of 1/k-symmetry modulation. Spatially tuned populations followed the same distribution, but only in variance (as predicted) and only for even symmetries, due to the 180° symmetry of integrated spatial segment activity from spatial tuning. Consequently, when these cell types are present, we expect the chance probability of having a dominant 6-symmetry to be 2nd or 3rd highest in the 4-9 symmetry window.

**Figure 2:**
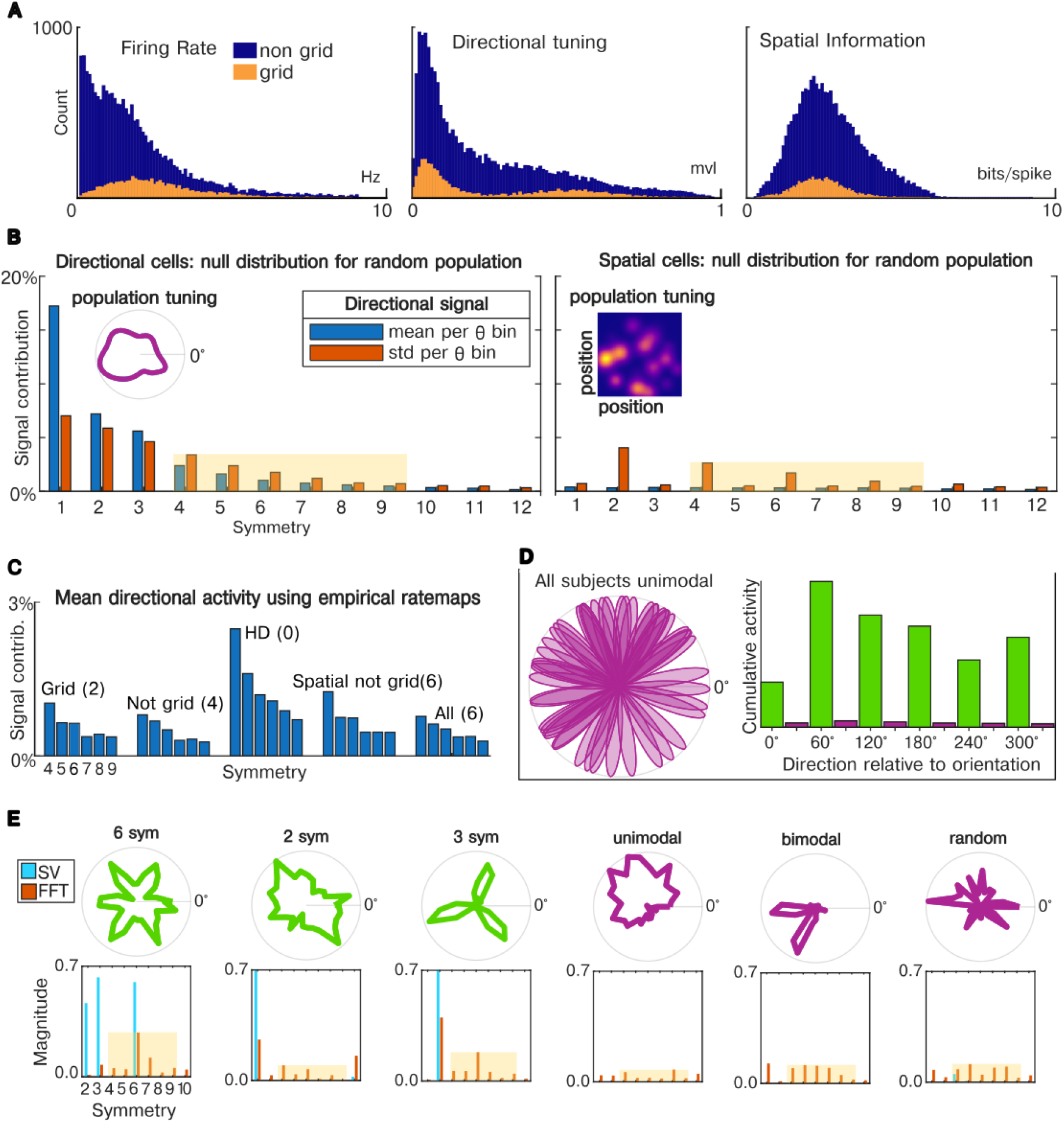
Null distribution of directional symmetry for directionally and spatially tuned cells. (A) Empirical rodent MEC neural populations contain many *non-grid* directionally and spatially informative cells. All cells in dataset pooled (Vollan 2025): n=23,453 across 29 open-field recordings^23^. Mean firing rate (Hertz), directional and spatial tuning specificity (mean vector length, spatial information). (B) Null distributions for random directionally-tuned (von Mises, mvl=[0.5-1], left) and spatially-tuned (2D Gaussians, right) cell populations (1-20 cells per sim), meaned over 100 Monte-Carlo simulations. We quantified symmetric modulation of mean and standard deviation of equal-length segments by direction using sinusoidal decomposition (see Methods). Signal contribution is normalized Fourier amplitude, expressed as a percentage. Inlay: examples of population level tunings. The subset of symmetries most often controlled against in hexadirectional analysis [4-9] are highlighted in yellow. (C) Empirical symmetry modulation of different cell populations. We combine empirical ratemaps with simulated trajectories (see Methods) for cell subpopulations from dataset (mean over n=29 recordings). Parentheses: n-recordings where 6-symmetry had highest signal. We show only the typical control symmetries [4-9]. (D) Induction of apparent hexadirectional modulation from non-hexadirectional data. We produce a common population-level plot showing hexadirectional modulation from subjects with a unidirectional response at random directions. For x-axis normalization, movement direction is shifted by ‘mean grid orientation’ (i.e. 6th harmonic phase). Per-subject, this ‘correction’ simply shifts the activity in 360° space so the peak aligns to the nearest multiple of 60°, producing an apparent 6-symmetric modulation at population-level. In sinusoidal decomposition phase is largely determined by location of peaks in input, such that this occurs for most non-uniform data. (E) Two methods for quantifying k-fold rotational symmetry. Fourier (FFT, red), which decomposes signal into sinusoids, and our method, Symmetric Variance (SV, blue), which computes cross-group variance after grouping by symmetry (see Methods). Top plot: input signal (green: rotationally symmetric inputs). Signal sampled over 30 direction bins. Gaussian noise of σ=0.2*Root-Mean-Square(signal) added. Typical control symmetries in hexadirectional literature [4-9] highlighted.

We performed this analysis on empirical sub-populations ^23^ (23,453 cells, 2A, 3B) in figure 2C for mean directional activity, per canonical hexadirectional analysis, using the hybrid method described (see Methods). On average, pure grid cells produced neither the highest amount of 6-symmetric modulation, nor the highest number of recordings where 6-symmetry was highest (in parentheses). Overall, these resembled a 1/f noise distribution.

This also highlights the critical need to approach this problem from a *spectral analysis perspective*. Figure 2D shows a common hexadirectional plot that can be spurious (see legend), which becomes clear if it is generated for multiple symmetries. We also propose an alternate measure of symmetry, *Symmetric Variance*, Figure 2E, which addresses some of the pitfalls of using sinusoidal decomposition as a proxy for k-fold rotational symmetry (see Methods).

### Conjunctive grid x head direction cells do not produce a hexadirectional signal

As discussed in the previous section, a spatial grid pattern alone cannot produce a directional modulation in mean firing. For this a second-order phenomenon is required. The second hexadirectional hypothesis is that unidirectional tuning of conjunctive grid-×-head-direction (grid-HD) cells cluster to grid axes such that the population level firing is hexadirectional. This was reported in the original hexadirectional paper^7^ for n=18 conjunctive cells across n=8 rats, but has not been replicated as far as we know. A replication was attempted in a medical thesis project^25^ in the Moser lab based on a dataset of n=224 conjunctive grid cells across 15 animals^26^, which was unsuccessful.

We also attempted to replicate this finding with a dataset of 29 MEC open-field recordings^23^ comprising a total of 23453 cells, 3699 of which were grid cells (see Methods). Plotting the directional tuning as mean vector length (MVL), we see a striking bimodal distribution between pure and conjunctive grid cells, figure 3B. Using the trough between peaks as our conjunctivity threshold (n=1349 conjunctive cells), we analyzed the relationship between directional tuning angle and grid orientation in a series of histograms, figure 3C. While there were clear groupings in grid orientation as expected from grid modules derived from multiple animals (19) across 29 sessions, we saw no grouping for directional tuning, which appeared flat (uniform) in both 360° and 60° space. To test whether head direction aligns with directional tuning grid orientation we quantified the circular distance between them in 60° space (3C rightmost). If aligned we would expect a peak difference at 0°. A circular V-test of this yielded pvalue=0.9 and v=-31 indicating no significant relationship. However, for the possibility that there is an offset other than 0° we ran a Rayleigh test for non uniformity, which had pvalue=0.03 rejecting the null hypothesis. The corresponding mean offset from the grid axes was 20°, however the MVL of this peak was very broad 0.05, almost flat.

**Figure 3:**
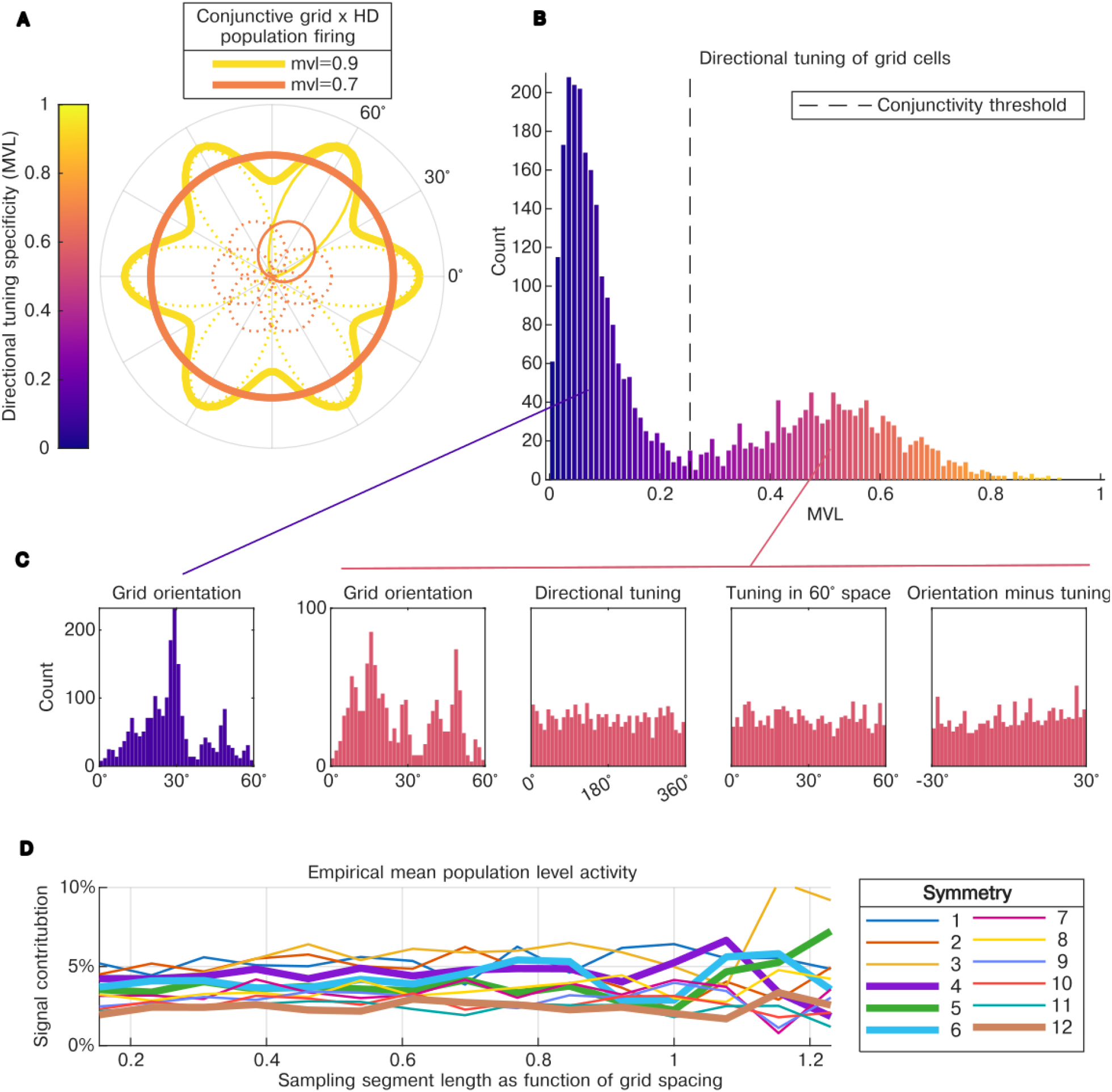
Directional signal of conjunctive grid × head direction cells. (A) Simulation of two conjunctive grid × head direction (grid-HD) populations with different tuning specificity (mean vector length, MVL). Each population consists of 6 cells with HD tuning aligned to one of 6 symmetric grid axis directions. All cells (thus both populations) have same total firing. Individual cell firing (modeled as von Mises) is shown in the thin/dotted line, combined population firing is shown with thick lines. R-axis = firing. At MVL=0.7 directional modulation is largely eliminated. Colorbar (panels A, B, C): directional tuning (MVL). (B) Histogram of grid cell directional tuning in dataset. Grid cells pooled across all recordings (recordings=29, grid=3699, pure-grid=2350, conjunctive=1349). We see clear bimodal distribution between Pure and grid-HD cells.We also see vast majority of conjunctive cells are below MVL=0.7 threshold required for directional modulation. Y-axis = count. (C) Grid orientation (all recordings) pure grid cells (purple), conjunctive grid-HD cells (red). For conjunctive cells, relationship between grid orientation and directional tuning is also shown. While grid orientation appears clustered (different modules, animals, recordings), also note difference between conjunctive and regular cells in grid-orientation distributions. Directional tuning was seemingly uniform (both in standard, and 60°space i.e. *mod*(*tuning*, 60°)). Rightmost panel: HD tuning does not align with grid axes (V-test p=0.9, V=-31), however the distribution of grid orientations minus HD tuning was not uniform (Rayleigh test p=0.03, but only slightly: MVL=0.05). (D) Empirical mean population activity (averaged over recordings), sample segment length expressed as ratio to mean grid spacing for recording (Ex: segment length = 80 cm, mean grid spacing = 40 cm, relative length = 2). If hexadirectional conjunctive tuning was present we would expect to see a high 6 symmetry regardless of segment length. Note rats change direction frequently so n-samples decreases significantly as segment length increases.

To scrutinize whether there was a session-specific offset between head direction tuning and grid axis we ran the previous analysis on each recording individually. The mean number of conjunctive cells per recording was 47, median=15 (∼5% of population), with 5 recordings having n>100 [104,116,119,130,173] (∼ 10% of population). For a V-test of mean tuning/grid axis difference of 0°, all but 1/29 recordings did not reject the null hypothesis (α=0.05). However, the recording that did pass had the highest number of conjunctive cells. 2 of 29 recordings rejected the Rayleigh test for uniformity at α=0.05 (MVL=0.2 for both, offset=[0°, 25°]). Notably, one was again the highest number of conjunctive cells, and the other had the highest percentage of grid cells (pure + conjunctive), 38% (next highest being 27%). Overall it appeared that there was no stable relationship between grid orientation and head direction tuning, however it was compelling that the recording with the highest number of conjunctive cells by approximately a 50% margin from next highest, did show a statistically significant Rayleigh-test result. That said, the recording with the next highest number of conjunctive cells, 130, same percentage of population, 16%, had a V-test p-value of 0.8.

There is a second, subtler requirement for this hypothesis: tuning specificity. Figure 3A shows an illustrative conjunctive cell population (n=6), with an ideal hexadirectional tuning distribution, for two tuning specificities. At fairly high tuning specificity, MVL=0.7, 6-symmetric modulation, though technically present, is negligible. Given the empirical directional specificity was MVL=0.51±0.06, it seems unlikely that conjunctive cells could generate a detectable hexadirectional signal. The recording with the highest number of conjunctive cells mentioned in the previously had a mean MVL=0.5, as did the next 4 highest.

Lastly, we examined empirical population-level mean activity in Figure 3D and computed directional symmetry by segment length. Unlike a grid spatial-tuning-response based hexadirectional signal, the grid-HD signal should not be modulated by segment length. Although 6-fold symmetry fluctuates, especially at longer segment lengths (most likely from sample sparsity, see next section), these fluctuations appeared noise-like and did not differ meaningfully from the 1/f null (Figure 2B-left). Here we show mean modulation across recordings (for per-recording plotting see Figure 4B ‘cell mean’, equivalent to population signal i.e. sum of cell activity).

**Figure 4:**
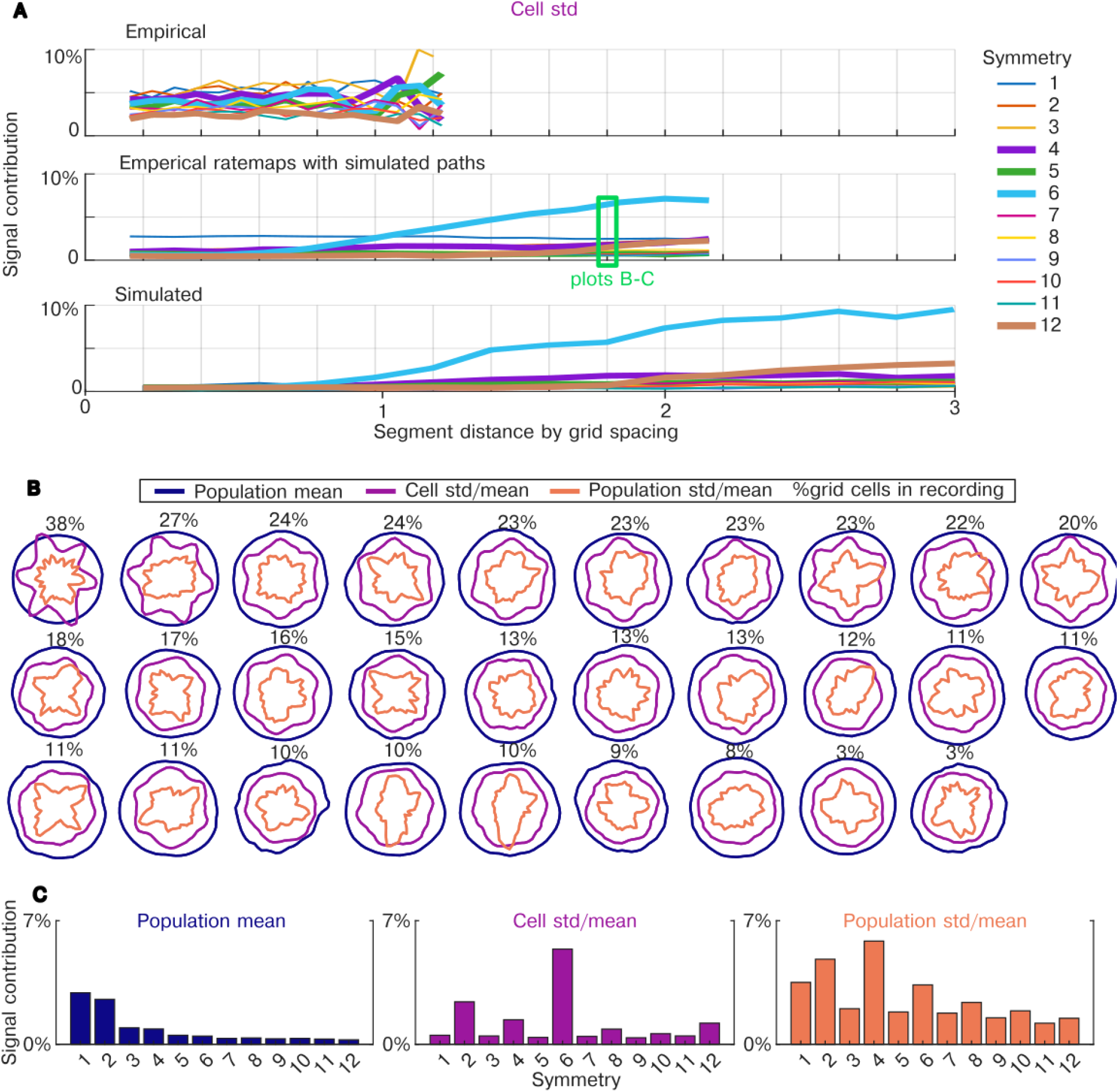
Directional symmetry in real and simulated MEC populations (cell, population, mean, variance) (A) Symmetry modulation in 23,453 MEC cells across 29 open-field recordings^23^. These empirical data contained a varying number of grid cells as well as spatially and directionally tuned cells, as would be expected in macro-level neural recordings such as BOLD. Due to grid pattern geometry, sample segment length affects symmetry modulation (discussed previously). Segment length is normalized by mean grid spacing per recording. We used three approaches to quantify cell activity: **Empirical (B top):** empirical trajectories directionally segmented and corresponding cell firing. While most parsimonious, rodents rarely run in long straight segments (excluding borders) yielding few samples at lengths where symmetric modulation is expected. **Hybrid (B middle):** empirical spatial and directional ratemaps and interpolating new trajectories. Green-highlight: ***data used for panels D-F***, 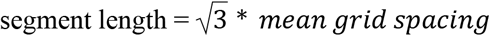 (theoretical length of maximum hexadirectional effect, Figure 1E). **Simulation (B bottom):** computational simulation tuned to match hybrid results, extrapolated for longer segment lengths. Mixed populations of n=1000: grid, grid-HD, HD, spatially-informative (2D-Gaussian), and noise cells at fixed proportions. Each length simulated 100 times, mean plotted. Cell-type proportions were optimized to match hybrid data rather than the empirical proportions, though they ended up being similar (see text). Notably, at n=1000 we could not reproduce the unidirectional signal contribution (sym = 1) seen in hybrid interpolation. More relevant symmetries plotted with thicker lines. “Signal Contribution” = normalized Fourier amplitude of directional signal, signal=standard deviation segment activity per direction. Polar sinusoid decomposition used as proxy for k-fold directional symmetry as per canonical hexadirectional analysis. (B) All cells in dataset pooled. Mean firing rate (Hertz), directional and spatial tuning specificity (mean vector length, spatial information). (C) Dataset plotted for 3 directional signals (calculated from all segments, per direction): mean population-level activity (blue), mean cell-level coefficient of variance (^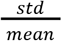^ i.e. CV, purple), mean population-level CV (orange), % - grid cells in recording. For display, signals are scaled to fit concentrically on plot (scales = [1.3x, 1x, 0.6x]). Recordings contain ∼20-350 grid cells. (D) Same data as D, directional symmetry modulation (signal contribution), averaged over recordings.

### The nonlinearity hypothesis

Next, we tested the nonlinearity hypothesis, which proposes that a nonlinear transformation of grid cell activity produces a hexadirectional signal. The nonlinearity hypothesis works by transforming spatial tuning derived directionally modulated variance into directional modulation mean, explaining the hexadirectional signal reported in the latter. To have an effect this nonlinearity must be applied to the integrated firing over a path (i.e. over time), not pointwise. Our aim is not to speculate in detail about any particular nonlinear physiological processes(es) in the transformation of cell-level action potentials to macro-level brain recording. Our goal is to address this from a theoretical perspective, constraining the description of a viable nonlinearity and estimating a reasonable upper bound on the magnitude of the hypothetical signal. Consequently, we consider two basic nonlinear transformations: power functions and thresholds applied discretely to total activity per segment.

#### The most theoretically likely type of nonlinearity

To produce the strongest hexadirectional response the nonlinearity is most effective at the cell-level (Figure 5A). This is because of the principle established previously that the grid pattern is most high contrast at the single-cell level and so has the highest directional variance. In contrast, the population-level pattern is close to uniform due to uniform distribution of grid phase (Figure 2D).

**Figure 5:**
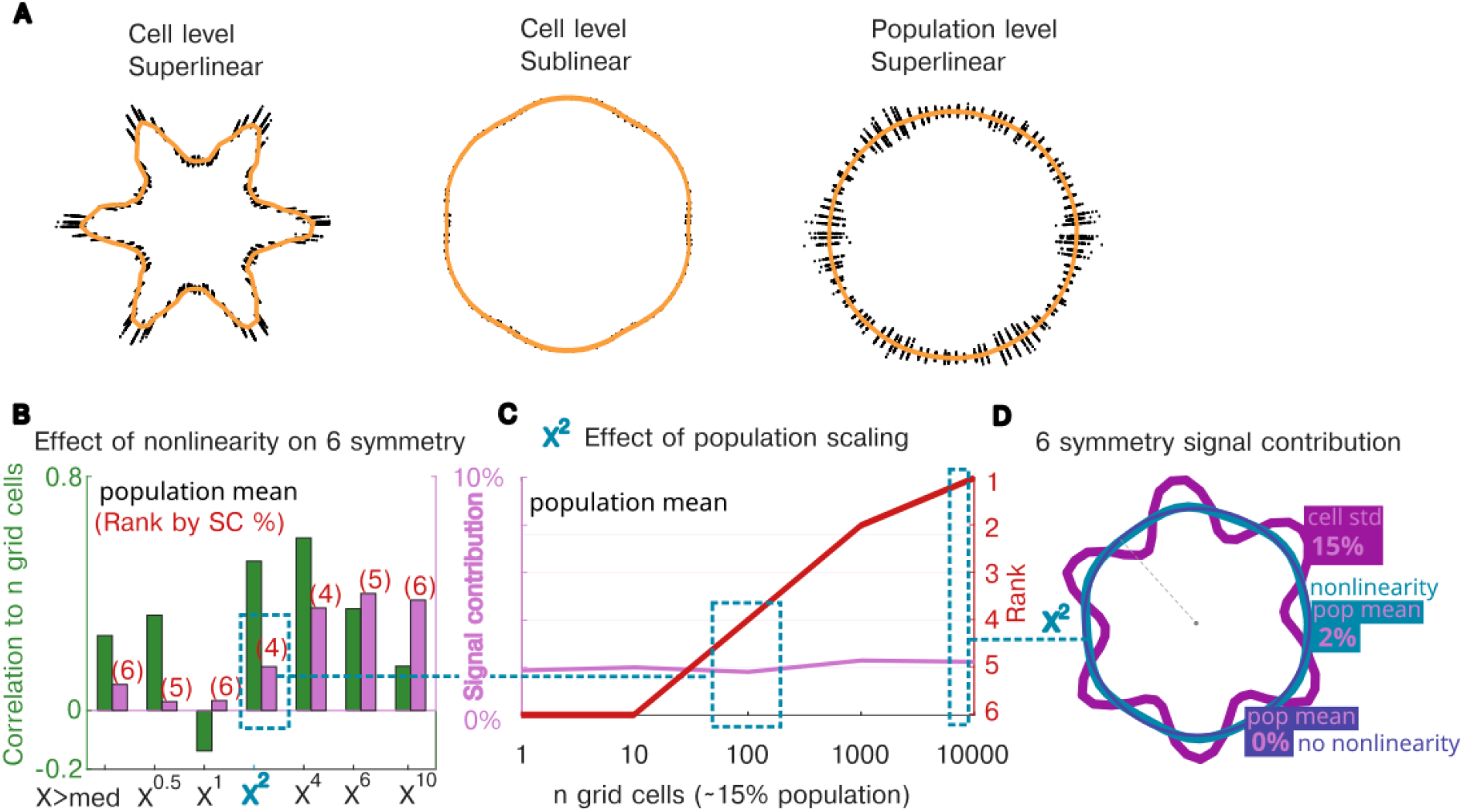
Hexadirectional modulation of nonlinearly transformed grid cell activity. (A) The effect of different nonlinearities on macroscopic grid cell activity. Plots show individual path segment (black points), θ-axis = movement direction, r-axis = population activity after nonlinear transform, mean in yellow. For each trajectory segment, we applied a nonlinearity *x*^*c*^ to total firing, for *c* = 3 (superlinear/expansive), and 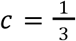 (sublinear/compressive). Key is *where* ‘*physiologically’* the nonlinearity is applied: to each cell before summing population (cell-level), or to summed population activity (population-level). A nonlinearity converts the uniform directional mean into a hexadirectional signal due to hexadirectional-variance produced by the grid tuning pattern (assuming trajectory constraints discussed). Modulation magnitude differs, with the largest signal produced by a cell-level, superlinear nonlinearity. (B) Different nonlinearities applied at per segment cell-level activity, where maximum effect is expected. We quantified population-level mean activity (per canonical hexadirectional analysis) for 6-symmetry only. Left y-axis (green): correlation of signal contribution to proportion of grid cells across recordings. Meaningful 6-symmetric modulation should correlate to proportion of grid cells in recording. Right-y-axis (pink) signal contribution meaned over recordings. Red parentheses: rank of 6 symmetry by signal contribution, rank-1st = highest magnitude, over all symmetries (excluding 0 i.e. mean response). Note, regardless of nonlinearity 6-symmetry did not dominate, despite dominant 6-symmetry in cell-level variance. (C) Scaling effect of population size on rank. We simulated a heterogeneous MEC population parameterized to match the signal contribution and rank from the hybrid approach (F) for an *x*^2^ nonlinearity, with increasing population size. This again approximately matched the mean empirical proportion of grid cells ∼15%. While hexadirectional modulation stays constant at ∼2% of total activity, the other symmetry contributions decrease such that rank of 6-symmetry increases. Blue dashed-line = hybrid data that simulation parameters were fit to (population size, signal strength, rank). (D) A visualization of post-nonlinearity population level mean signal contribution (2%, teal) vs. the cell-level standard-deviation (15%, purple), and no nonlinearity (blue), for the maximum population size (10,000 grid cells, ∼15 of population).

Similarly, relative to the scale of the original activity, a *superlinear* (expansive) physiological process (facilitation, gain enhancement, potentiation, etc) will produce a stronger modulation than a *sublinear* (compressive) process (adaptation, repetition suppression, saturation, etc), for the same scale of measurement. This is intuited by comparing two values greater than 1 after squaring versus after rooting. We expect to see a larger difference between the values after squaring. We simulated these principles in figure 5A for a grid cell population. We see that a superlinear transformation of firing at cell-level caused the highest symmetric modulation in the mean (yellow). A sublinear nonlinearity at cell-level produced a visible, but weaker modulation due to its compressive nature (note also the signal is at the off-axes, i.e. ‘misaligned’). A nonlinearity at the population level, even a superlinear one, produced virtually no modulation relative to the overall signal magnitude.

#### Quantifying prerequisite hexadirectional variance empirically

Working off this premise, we focused on cell-level nonlinearities. As mentioned, a prerequisite of this hypothesis is hexadirectional variance. To begin, we verified that this variance pattern exists empirically. As discussed the spatial-tuning hexadirectional signal is caused by the varying distance required to cross consecutive firing fields in different directions. Detecting this requires having straight trajectory samples of lengths at least close to grid-spacing. A central problem in investigating the hexadirectional signal with rodents is that in open environments they rarely follow long straight paths except along borders. Critically, *this alone* could explain why the hexadirectional signal has not been reported in rodents. We nonetheless analyzed the variance modulation directly in our rodent data (Figure 4A top), which did not show 6-symmetric variance modulation, even at segment lengths predicted in simulation (4A bottom). However, we could not preclude that this was due to decreasing number of samples at longer lengths.

#### Hybrid approach

Despite continually adding parameters to our computational model including: elliptical grids, field location noise, etc, (see Methods), we could not replicate the irregular empirical distributions of spatial and directional cell-tuning (Figure 2A). To remedy this, as well as the lack of longer segment samples, we used a simple hybrid approach for bootstrapping new data: treating the empirical spatial and directional ratemaps of each cell as joint independent marginals, and sampling new segments (see Methods). Note: these segment lengths were still constrained by the real arena dimensions relative to the empirical grid spacing.

Figure 4A shows a Fourier decomposition of directional firing for three sources of data: empirical, hybrid, and simulated. We measured the mean cell-level standard deviation over segment activity per direction, as we expected the highest hexadirectional signal. We used standard deviation, not variance, to stay in the same relative scale as the original signal, as this is what will be input into any downstream nonlinear process. The mean over recordings is shown. From these plots we estimate that a sampling segment length of at least ∼1.3 times the mean grid spacing is needed for a reliably dominant 6-symmetric signal (under the assumptions of this model). For the hybrid data this modulation was around 6% of the total standard deviation of the signal. According to the simulation this could increase up to 10% at greater lengths. Interestingly we did not see the theoretical signal peak at *sqrt(3)*grid-spacing* as in figure 2E, this is likely due to noise from having non-grid cells in the population reducing overall signal, (note difference y-axis limits). That said we do see that the 12-symmetry increasing in magnitude. To compare other aspects of the signal, we fix segment length at *sqrt(3)*grid-spacing*. Besides the reasons mentioned, this length was obtainable in most recordings, and was used for the remaining analyses.

To better understand the distribution of symmetric modulation in this dataset we have plotted 3 directional signals for each recording individually (4B), with mean plots in 4C. We use the coefficient of variance, CV, 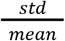, to account for mean fluctuations that are theoretically unrelated to spatially firing (as discussed), and to maintain relative scale to overall signal magnitude, important when considering detectability. Cell-level CV (purple, 4B) demonstrated clear correlation between grid cell proportion and hexadirectionality, suggesting a causal relationship. At the aggregate level, we observed a dominant 6-symmetry at ∼5% of the total CV (4C), despite the empirical noise of a mixed directionally/spatially-tuned cell population (2A). Though trajectories are extrapolated, *this is the most empirical evidence of a rodent hexadirectional signal that we are aware of*. As predicted, mean activity (blue) has little directional modulation, and generally resembles the 1/f null distribution of random directional signals (Figure 2B-left, Supplementary Figure S1A). Both cell and population level variance (4C, purple/orange) show the hallmark of spatial-tuning-derived directional activity where even symmetries are elevated (2B-right). Population-level variance resembles a null spatial tuning distribution though 4 and 2 symmetries are inverted. However, when viewed individually (4B, orange), the population-level variance signal had unexpected structure; not hexadirectional, but clearly modulated by direction: seemingly aligned to grid axes at times, and orthogonal at others, with the highest symmetry being 4-fold, in contrast to the null expectation.

#### Applying a nonlinearity

Given we detected hexadirectional modulation in cell-level variance, we explored some basic nonlinearities (power functions, thresholding) on cell-level activity (5B, 5C). We quantified only 6-symmetry 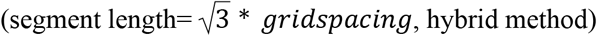. Maximum correlation of modulation to grid cell proportion (left-axis, green, across-recordings) occurs at *x*^4^; with minimum at *x*^1^ (no transform/nonlinearity) as expected. Comparing magnitude (right y-axis) we see max of 5% (*x*^4^, *x*^6^, *x*^10^). In red we quantify rank by magnitude over all symmetries (1^st^=highest i.e. dominant), which more clearly contrasts against the null distribution. We see peak rank=4^th^ for *x*^2^, *x*^4^. Consequently we consider *x*^2^ the weakest nonlinearity to produce the highest signal, based on raw magnitude. 6-symmetry rank=4^th^, while not dominant, is higher than chance for non-grid directional populations (6^th^), though lower than chance for random spatial ones (3^rd^) (Figure 2B, Figure S1A-rightmost). *x*^2^ has significance in nonlinear natural processes as *a*_2_ *x*^2^ is the lowest-order nonlinear correction to a linear model (*a* _0_+ *a*_1_ *x*). We focus on *x*^2^ in Figure 5C. The sublinear (compressive) transform we tested, *x*^0.5^ appeared not to produce a detectable signal. The thresholding nonlinearity, *x*←{0, *x* < *median*(*x*)}, was also non-viable. Higher thresholds may work, but may also be susceptible to noise similar to the high exponents seen here.

#### Scaling effect

Applying basic nonlinearities did not produce strong evidence for detectable hexadirectionality in macroscale signals like fMRI-BOLD: 0-5% modulation, rank close to chance, in the ground-truth neural activity. However, a decisive scaling factor exists not present in rodent data. For smaller MEC populations, sparse amounts of directionally, or spatially tuned cells produce anisotropical activity that is more disruptive to symmetric modulation. In larger populations with the same cell-type proportions, the directional and spatial noise from non-grid cells ostensably becomes uniformly distributed. Though our empirical data contains populations of *n* ≈ 1000 cells, we could simulate up to *n* ≈ 1 *million* cells (see Methods). To test scaling, we parameterized our simulations to produce the mean signal curve of the hybrid approach and increased the population size (Figure 5C). We see that while hexadirectional signal magnitude remained low at 2% (Figure 5D), its rank increased. Though seemingly contradictory, this signifies that while 6-symmetry magnitude relative to overall mean signal remained the same, the noise-based symmetries decreased. Importantly, this effect relies on uniformly distributed spatial and directional cell tuning. Exploring the scaling effect over power function exponents, unexpected optimality existed for exponents between [*x*^1.5^, *x*^3^], with signal strength peaking at around 2% and dominant symmetry occurring at ∼10000 grid cells Figure S2.

## Discussion

We tested the hypothesis that grid cells produce a hexadirectional signal observable at the macro/meso-scale (fMRI, MEG, iEEG). Despite the widespread application of hexadirectional analysis, a clear account of how grid cells produce this signal has been lacking. To address these gaps, we evaluated the three leading hypotheses^21,22^ using largescale rodent grid cell recordings^23^ (23,453 MEC cells across 29 recordings), and computational simulations, as well as the analytic method itself. A recurring theme is that the answer depends on scale: single grid cells robustly carry sixfold directional structure, but in their variance rather than their mean, and pooling cells of different spatial phase largely erases it unless a nonlinearity acts at the cell level before pooling (Figure 1C–D, 5A).

We have shown that based on first principles and empirical data, grid cell firing alone does not produce a hexadirectional signal in mean directional response: the quantity measured in the literature. This is true for any spatially modulated firing pattern by construction. This invariance assumes that space is sampled evenly across directions; uneven sampling can modulate the mean, but then the modulation reflects the trajectory rather than the grid. What grid geometry does predict is a sixfold modulation of cell-level variance, which we observed in cell-level activity derived from empirical ratemaps. Hexadirectionality in mean cell-level variance scaled with the proportion of grid cells in a recording (Figure 4B–C); consistently, the mean directional activity of pure grid cells was no more sixfold than that of other MEC cell types (Figure 2C). In terms of hypotheses relying on secondary phenomena to produce a *mean* hexadirectional signal, In the 1,349 conjunctive cells examined here we found marginal evidence that the directional tuning of conjunctive grid cells is anchored to the grid axes, the low specificity of this tuning argues against this explanation as a general mechanism: alignment in particular populations cannot be excluded (the recording with the most conjunctive cells showed it and the rest did not; Figure 3C), but even perfectly aligned cells with the observed tuning width (MVL ≈ 0.5) produce negligible population modulation (Figure 3A).

While a number of non-linearities may underlie the neurovascular coupling of the fMRI-BOLD signal^27–31^ this remains an active field of research^30,32–34^. We have shown, theoretically that the nonlinearity hypothesis is viable and have identified its most likely physiological description: a cell-level, superlinear (expansive), ∼*x*^2^ transformation of path segment firing. This nonlinearity must act on activity integrated over a segment (an instantaneous nonlinearity has no effect), and before summation across cells: once cells of different spatial phase are summed, little directional variance remains for any nonlinearity to convert (Figure 5A). Stronger nonlinearities were not uniformly better: intermediate power-law exponents provided the best combination of contribution and rank, so maximizing amplitude alone did not identify the most specific signal (Figure 5B).

A nonlinearity combined with an increased number of grid cells contributing to the population signal (∼ 10k+), a hexadirectional signal emerged from the ground-truth, single-cell firing at a ∼2% sixfold signal contribution (sixth-harmonic amplitude as a fraction of the summed harmonic amplitudes including the mean; Methods) for a heterogeneous MEC-like population (Figure 5B-D). Increasing the number of grid cells did not raise this magnitude but made sixfold the dominant symmetry, because competing directional structure from non-grid cells averages out (Figure 5C). Regarding the number of neurons in the human EC estimates range between 5k-15k cells per mm^3^ depending on layer and location^35–37^. For a 3 × 3 × 3 voxel this would correspond to 135k-405k neurons. In rodents grid cells can exceed 30% of all recorded MEC cells^24^ (in our dataset max 38%, mean 16%, not including interneurons, Figure 4B). Thus their count could exceed 120k+ cells in such a voxel, which would support the scaling effect mentioned above, though this is using most favorable estimates and assuming appropriate sampling of the pmEC (human homologue of rodent MEC with a volume of ca. 600mm^3^)^38–41^. Critically this assumes that grid-cell fractions from targeted recordings hold across the sampled tissue, whereas increasing evidence points to functional cell types cluster anatomically within MEC^24^. Consequently, even an ROI restricted to MEC, which already exceeds the anatomical specificity of many human measurements, could include enough non-grid tissue to dilute the grid-cell fraction below that required to produce the signal.

There are other reasons to believe this ∼2% could be inflated. In addition to selecting a segment length optimized to grid-spacing (Figure 1E right, 4A), we apply a nonlinearity to the neural activity of each segment and direction individually; there is no continuity in time. While mathematically expedient, this implies a precise, directionally-gated physiological mechanism. Alternatively, our results would reflect a time-based nonlinearity for trajectory segments matching the time constant. Varying speed and segment length adds significant noise to an arguably tenuous hexadirectional signal. This magnitude relies on an expansive, superlinear transform, which are less often hypothesized in this context compared to compressive, sublinear ones (adaptation, repetition suppression). Moreover, the nonlinearities described for neurovascular coupling may only act on aggregate activity, i.e. after pooling, whereas the required transformation must act per cell before pooling; expansive processes at that level (facilitation, gain enhancement, potentiation) exist but have not been linked to this signal. In addition this dataset was collected to maximizing the number of grid cells and postprocessed to exclude fast-spiking putative interneurons, likely inflating our population proportions. We did not systematically address the impact of different sources of noise, such as physiological, analysis or measurement related, as it goes beyond the scope of our proof-of-concept approach. Our estimate therefore upperbounds the expected size of the signal with single cell precision, rather than its detectability at the meso/macro-level per se, which also depends on measurement noise and trial numbers. In the same vein, we did not include human fMRI data, because we assumed a ground-truth, single cell perspective.

In doing hexadirectional analysis, two questions must be kept apart when interpreting the measured signal: whether it reflects rotational symmetry at all, and whether sixfold structure is expressed preferentially over other symmetries.

In considering the null hypothesis, we have highlighted pitfalls in hexadirectional analysis (see Methods). Namely, conflating peaky directional activity with symmetric activity, and computing a single harmonic decomposition without contextualizing it systematically using spectral analysis. A hexadirectional analysis not comparing other harmonic components essentially can only prove the presence of stable directional peaks. A robust spectral comparison should be the central analytic approach, not a marginal control method. In this regard, we have generated the null distribution of symmetric directional modulation for random populations of directionally and spatially tuned cell populations (Figure 2B). We have shown that in commonly used control symmetries, a dominant 6-symmetry is 3rd or 2nd highest by chance (note that in the case of spatial cells, a nonlinearity must be present to produce a modulation in the mean), and that pre-processing can affect false-positive statistical inference (Figure S1). Stronger evidence for preferential sixfold structure requires evaluating the sixth-harmonic response magnitude relative to an appropriate spectral null distribution. We have also shown that a popular and visually convincing, group-level plot of the hexadirectional effect easily results from random data (Figure 2D). We have also described a new measure, Symmetric Variance, that better captures rotational symmetry independent of peak shape.

The MEC is also home to other directionally tuned cells^2,24^ (Figure 2A). Unlike pure grid cells these cells are defined by the strong peaks they produce in directional firing. Consequently, these neurons will have large impact on directional analysis. This holds particularly where their preferred directions are anchored to features that are stable within a session, such as landmarks, boundaries or goal locations. We have shown how sinusoidal decomposition is not an ideal proxy for measuring rotational symmetry and can yield unreliable symmetry estimates (Methods, Figure 2D, Figure S1). Specifically, we showed that directional activity peaks produced by cell-types such as these produce high, temporally stable measures of symmetric modulation (Figure 2B, Figure 2D, Figure S1D, Methods).

A compelling case for the hexadirectional signal is the replicated observation that it correlates with behavioural performance^7,42,43^. If in fact stable non-symmetric directional activity from non-grid cells are responsible for the signal reported in these studies, it would still explain the correlation between navigation circuitry and cognitive performance. As shown, this non-symmetric but stable directional activity would have a reasonably high chance of passing controls in hexadirectional analysis (partitioning data, non-zero mean, and naive shuffling^44^, Figure 2B, Figure 2E, Figure S1A, Figure S1D, Methods).

A major difference between human grid cell research and the rest of the grid cell field is that the effects in hexadirectionality are commonly only statistically tested at the group level, with simultaneously very few clear single subject examples of the hexadirectional signal. Arguably this indicates that hexadirectional analysis is currently not reliable enough for subject-level clinical applications. Recent studies of the human entorhinal cortex using precision fMRI especially highlight the challenge of translating group-level findings to individual-subject level^45,46^ Another idiosyncrasy is that positive results often only occur after dividing the data into specific subsets: e.g. by behavioral state, task epoch, participant group, or (sub)frequency band. If these subdivisions were standard across studies this would not be an issue, but collectively, these post-hoc subdivisions inflate the Type-I error rate. In simulation, allowing analysis-pipeline flexibility across hemisphere (3 choices), epoch (3), and ROI (2) yielded a significant sixth harmonic with nonsignificant controls in roughly half of null cohorts of 20 participants (Figure S3). Such flexibility need not involve deliberate selection^47^. Our analyses quantify this risk; they do not establish selection in any particular study. These issues have been discussed in the broader context of the replication crisis^48,49^.

Complementary computational work^22^ addresses these same three hypotheses. Broadly speaking, for the geometry hypothesis, which we argue is non-viable, they find hexadirectional modulation, but state it is either spurious, or dependent on starting location. They find an effect of similar magnitude, but more stable, with nonlinearity (adaptation/repetition suppression), though this is with favorable decay parameters, and constant speed/grid spacing/orientation. They perceive the conjunctive hypothesis as computationally and empirically strongest, however this is based on parameters from older, smaller studies^7,50–52^ that were not replicable in the larger dataset^23^ used here. They also describe a false positive caused by the hexadirectionality of the sampling path itself. It is noteworthy to mention the parallel between this line of work and grid cells in 1D linear track environments. In both contexts the 2D grid pattern is collapsed to 1D. In particular, previous work^53^ has described sampling the 2D grid pattern in straight paths at different directions. This work focused on single-cell-level activity in fixed directions, while we focus on the overall directional modulation at the population-level, we nonetheless arrive at many of the same findings. Just as importantly, we converged on the same methodology of using Fourier^53^spectral analysis.

To bolster the hexadirectional hypothesis going forward we suggest quantifying stable directional activity peaks as standalone effects, significant in their own right. For detecting grid cells non-invasively we also recommend measuring directional variance, though we predict a small signal at the population/macro-level (sixfold structure dominates cell-level, but not population-level, variance; Figure 4C). Notably, in this study we showed a hexadirectional variance pattern only with extrapolated trajectories. It would be ideal if hexadirectional variance were shown empirically with rodents running longer (straight) paths, as well as to test the nonlinearity hypothesis. Conversely, human virtual-navigation tasks supply the long straight runs that rodent foraging lacks and may be better suited to express a geometry-dependent signal, although the unknown spacing of human grids complicates matching path length to it. Alternatively, to support the physiological basis of the nonlinearity hypothesis, more research on neurovascular coupling is needed, especially cell-level coupling, as well as stronger computational modeling of the hypothesized signal. To consolidate observations across studies more standardization is needed^54^, as outlined. Simple, individual level plots with minimal pre-processing (direction vs activity) should ideally be reported in addition to group level second-order statistical plots to build a better intuition of the typical directional signal. The nonlinearity account is specific enough to be tested: it predicts a dominant sixfold variance signature in single grid cells on long straight paths; a mean signal that grows with the number of contributing grid cells; and a dependence on path length relative to grid spacing that a conjunctive-cell mechanism would not show (Figure 3D, 4A). These predictions distinguish a grid-cell source from stable non-grid directional tuning. Given the diversity in where and how the signal is observed, preregistering analysis pipelines is desirable. Perhaps most critically, with the missing body of clear single subject examples, the non-parsimonious nature of the nonlinearity required, the lack of its evidence, and the prediction of a weak hexadirectional modulation if it does exist (Figure 5D), a high likelihood of a significant 6-symmetric modulation of MEC activity should not be assumed.

Therefore, analyses showing a 6-symmetry should ideally do so also against all other symmetries as constrained by the sampling process, regardless of method.

In light of the ground truth MEC populations analyzed and methodological pitfalls described, evidence for hexadirectional signals outside the entorhinal cortex warrant heightened scrutiny. Such claims require particularly compelling evidence in brain regions lacking electrophysiological support for single-unit grid cells. Within the entorhinal cortex itself, a grid-cell source predicts stronger effects in the posteromedial subdivision (pmEC, proposed homologue of rodent MEC) than in the anterolateral subdivision (homologue of LEC, which carries little spatial signal)^38–41^ ; whole-EC analyses cannot test this, and comparing pmEC, alEC and extra-entorhinal regions at matched sensitivity would sharpen source attribution either way. Anatomical diversity of reported effects does not exclude a grid-cell contribution; it leaves open whether a common sixfold component has a common cellular source. Indeed, even in the rodent MEC (or human pmEC) - the region with the most robust evidence for grid cells - a mechanistic link between single-unit grid-cell activity and hexadirectional population signals has hitherto not been clearly demonstrated.

Here, we demonstrated critical limitations of hexadirectional analyses in detecting grid cell population activity, as well as the merit of this approach under different hypotheses. This work has further implications for bridging single-cell activity to population signals like fMRI-BOLD in the EC and beyond.

## Resource availability

### Lead contact

*Requests for further information and resources should be directed to lead contact, Noam Almog <u>noam</u>*.*<u>almog@ntnu</u>*.*<u>no</u>*.

### Materials availability

*NA*

### Data and code availability

*Available upon publication*

## Acknowledgments

We are grateful for helpful discussions with Yoram Burak, Jan-Mathijs Schoffelen, David Attwell, Noam Shemesh, Ila Fiete, Michael Hasselmo, Jeffrey S. Taube, Dori Derdikman, Martin Stemmler, Andreas Herz, Richard Kempter, Yasser Roudi, Ines Samengo, Patrick J. Drew, Ikhwan Bin Khalid and help from Natalie Ruth St. John.

We thank Abraham Vollan and Edvard and May-Britt Moser for providing the rodent data. The work was supported by two Research Council of Norway Centre of Excellence Grants (Centre of Neural Computation, Grant number 223262; Centre for Algorithms in the Cortex, Grant number 332640). In addition, the project was funded by the Kavli Foundation.

## Author contributions

NA, Conceptualization, Data curation, Software, Formal analysis, Investigation, Visualization, Methodology, Writing. TPD, Conceptualization, Methodology, Writing. TNS, Conceptualization, Resources, Supervision, Funding acquisition, Methodology, Writing.

## Declaration of interests

The authors declare no competing interests.

## Declaration of AI technologies in the writing process

The authors used ChatGPT, Claude, and Gemini for paraphrasing and grammatical corrections. The authors take full responsibility for the content of the publication.

## Supplemental Information

Supplementary.pdf: Figures S1–S3.

## Methods

### Vollan 2025 dataset

This dataset contains electrophysiological recordings from 19 freely behaving rats implanted with high-site-count Neuropixels probes in the medial entorhinal cortex (n=10), the hippocampus (n=2), or both regions (n=7). Numbers of simultaneously recorded units ranges from 235-1,522 and included all cell types except interneurons. For this study we did not use the hippocampus only recordings. Of the different navigational tasks, we only used the open field sessions for a total of 29 recordings (n=18 rats). These rats foraged for randomly scattered food crumbs (corn puffs or vanilla cream cookies) in a square open-field box with a floor size of 150×150 cm and a height of 50 cm. The floor was made of black rubber, and the walls were made of black expanded PVC plastic. Notably one of these openfield sessions occurred in a darker lit, circular environment 150 cm in diameter. This recording was not remarkable in any statistical way we observed 6th highest grid cell account, etc, so we did not exclude it from the dataset. The animals’ head positions and orientations were tracked with OptiTrack motion-capture system. The raw neural activity was spike-sorted using Kilosort 2.5. Cells with mean spike rate of less than 0.1 Hz or greater than 10 Hz were excluded. For a more detailed description see^23^.

### Quantifying rotational symmetry

Hexadirectional analysis has multiple implementations; here we focus on the most common approach: 6-period sinusoidal regression. Nonetheless, the majority of this discussion applies to other implementations such as Representational Symmetry Analysis (RSA).

#### Hexadirectional analysis disambiguation

The most common hexadirectional analysis method is a form of circular regression where the 6 harmonic modulation (60° periodicity) is calculated (Eq. 1 below). In terms of nomenclature this is technically multiple regression, and thus it is technically a General Linear Model (GLM) and also a Generalized Linear Model (also GLM).

In the context of hexadirectional analysis the phase of the 6 frequency component is dubbed ‘mean grid orientation’ and magnitude (amplitude) is ‘hexadirectional modulation.’

#### Canonical hexadirectional analysis and Fourier decomposition

In hexadirectional analysis, the amplitude of polar sinusoids are used as a measure of k-fold rotational symmetry in an angular signal, specifically for the 6th harmonic. This is computed using the following regression:

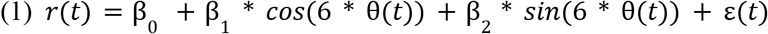

Which implies (under standard regression assumptions)

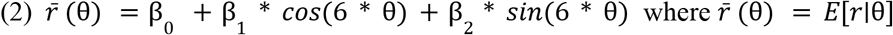

Such that

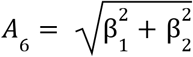

This is equivalent to the Fourier series for the 6th harmonic:

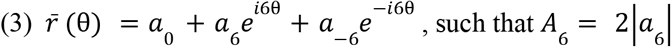

When angular sampling is uniform (Δθ, samples per θ), both approaches produce the same magnitude *A*_6_, and phase ϕ_6_.

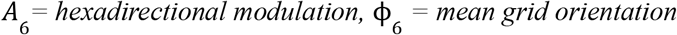

When directional sampling is uneven (more samples for a given direction and/or directions are not evenly spaced), outputs vary. In the latter case Fourier decomposition requires an explicit interpolation step.

Advantageously this explicit interpolation step forces the user to address irregular sampling biases explicitly, as opposed to regression where they can go unnoticed. Another advantage is computational efficiency of the Discrete Fourier Transform, especially when quantifying multiple symmetries, which are calculated simultaneously.

#### Importance of the spectral perspective

In canonical hexadirectional analysis the 6-period sinusoid regression is treated largely as a standalone model. Analyzing the 6th harmonic on its own is a major pitfall. The magnitude of any single harmonic largely reflects overall signal scale and is only meaningful relative to the remainder of the spectrum, and an appropriate null distribution. Natural signals contribute power across many harmonics. Thus the null expectation of this metric is not zero, but a nonzero, typically nonuniform distribution (Figure 2B, Figure S1A). Specifically the 1/f null distribution is common (f = k in k-symmetry context). Consequently, statistically testing harmonic magnitudes against zero is mis-specified.

Many hexadirectional studies do control for other harmonics/symmetries, however this is done idiosyncratically, typically for some subset of k = 4-9. However, comparing in only a narrow band provides incomplete spectral context which can distort overall inference (Figure S1C). This can be in Figure 2D (Fourier=orange) in which a 3-symmetric signal through a 4-9 band (yellow highlight) appears harmonically as a 6-symmetry.

A spectral perspective also elucidates essential constraints. Sampling-rate (angular bin spacing) and smoothing windows limit which symmetries are detectable, which is not necessarily apparent if not approaching this method from a spectral analysis perspective. Full spectrum decomposition also enables normalization (see below) and comparability across datasets, and is more interpretable than reporting an unbounded regression coefficient (β).

One notable methodological choice in this work is not to subtract the mean from our signal before fourier decomposition. The reason for this is that we are not using the decomposition purely to describe periodicities in our signal, our goal is more broadly to get a sense of how periodic activity at the micro scale is detected at the macro scale. If the entirety of the symmetric modulation comes down to x number of spikes, we want to know if the mean of the population activity is closer to 10x or 10000x i.e. how big of a modulation should we expect. While in theory not removing the mean can cause spectral leakage in the low symmetries (frequencies). Fortunately in our case, we care only about whole symmetries (6-fold not 6.1-fold), which are exact harmonics of our sampling window (i.e. 2pi/n where n is always an integer), in which case the fourier bases are entirely orthogonal, meaning they are not affected by leakage from the mean.

With the mean included we can more easily relate the magnitude of any given symmetry to the full spectrum of periodic activity in the signal as well as the overall magnitude of the signal itself. In our case we do a simple normalization of the magnitude (amplitude) of a particular symmetry divided by the total sum (including the 0 symmetry which is the signal mean) 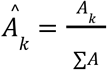 We then express this as a percentage and call it “Signal Contribution”. This gives us a sense of the modulation of the signal at a given periodicity relative to the magnitude of the entire signal. It could be argued that a better measure would be power 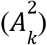 instead of magnitude. For statistical inference, the power would tell us how much each symmetry contributes to the variance of the angular signal (Persival’s theorem), however this is only in the case if we subtract the mean. Since we are not subtracting the mean, we chose the former measure for the reasons described. Importantly, the choice of using the coefficient magnitude (*A*_*k*_) also keeps symmetry modulation in the scale of the actual signal (similar to the standard deviation), without squaring the magnitudes (as in the variance), which is easier to intuit. That said, we have observed anecdotally that power, due to the nonlinearity of squaring, can do a better job of dampening noise symmetric modulation when values are normalized between [0 1] as above.

All that said, sinusoidal decomposition is an imperfect metric for rotational symmetry.

#### Quantifying rotational symmetry with sinusoids

There is an implicit bias in using sinusoid amplitude as a proxy for k-fold rotational-symmetry, which is that the signal is k-periodic sinusoid shaped (square-wave shaped in the case of RSA). This may not be the case, for example when the hexadirectional signal is generated by grid x HD cells with narrow directional tuning. In general, a perfectly k-fold symmetric pattern will have different power in the decomposed k-frequency depending on how well the signal fits a sinusoid. Conversely, the decomposition of an asymmetric signal will have a spectral profile dependent on how well the widths of its peaks fit a k-periodic sinusoid. In other words, the k-component projection of a sinusoid can be driven by one or several strong peaks whose angular positions and/or shape happen to align with that sinusoid. In this case, the signal reflects directional tuning with incidental harmonic structure, not k-symmetric peaks spanning over 360° (Figure S1D). Demonstrating symmetric activity should require evidence of multiple peaks with approximately k-fold spacing rather than isolated features that merely ‘resonate’ with a sinusoid.

#### Symmetric variance

To more parsimoniously measure symmetry in a directional signal, *S*_θ_, we introduce Symmetric Variance (SV), which quantifies how much variance is explained by symmetric grouping, (Figure 2D). This simple, normalized measure works by comparing the fraction of signal variance explained by across-group-variance, when the signal is grouped by coordinates that are equivalent according to the symmetry. While it can be implemented for any symmetry; in this context we use k-fold rotational symmetry.

To quantify the amount of k-fold symmetry in directional signal *S*_θ_, we group the values such that:

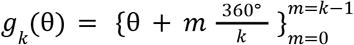

For example to test 3-fold symmetry, we would group the directions [1°,121°, 241°] together.

We then take the mean of each group, and divide the variance across groups by the total signal variance:

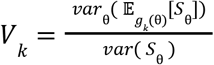

We then correct this value by the expected reduction of variance, 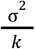 if the values in the groups are independent:

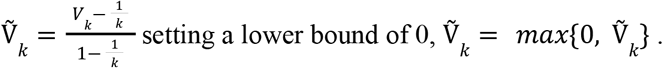

Lastly, as an error checking step we compare subsymmetries. For example if a signal is 6-fold symmetric, it must, by definition also be 2- and 3-fold symmetric. To be conservative, we assign the value to the symmetry as the minimum value calculated for the symmetry its sub-symmetries:

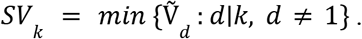

The result is a unitless metric between [0 1] that is invariant to phase and agnostic to signal shape. It can be viewed as a simple ANOVA-style variance decomposition, that quantifies the fraction of total angular variance explained by between-group-variance (grouped by symmetry in this case), directly aligning the method with standard variance-based effect size and reliability measures (e.g., ANOVA-η^2^ or Intraclass Correlation Coefficient).

In simulation this method appears more robust to noise, and the artifacts related to signal shape mentioned above (Figure 2D).

For a more conservative metric, 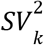, is even more robust to noise. Another advantage of this metric is that since the computation for symmetry with Fourier is fairly different, the two metrics yield fairly different results when the signal is noisy. Consequently another useful metric is *A*_*k*_ * *SV*_*k*_, especially when done for the whole spectrum. While not perfect, it is arguably more parsimonious, at the very least compliments the sinusoid approach when analyzing noisy signals.

#### Regression betas in hexadirectional analysis

A central measure of hexadirectional analyses in the literature is the contrast of beta estimates of the aligned and misaligned conditions^42^, or alternatively, the contrast of the beta estimate of an aligned cosine regressor and baseline^7^. Beta parameter estimates in GLMs can be signed and are unbounded. Negative beta values are not uncommon for the aligned condition for individual participants. However, it is arguably unintuitive how to interpret these negative values (which imply that the preferred mean grid orientation has changed orthogonally between partitions). Instead, a more intuitive approach would be to report the actual preferred angles (the phase of the Fourier decomposition, ϕ_*k*_) of the different data partitions (see for example^42^), as well as the signal magnitudes, because these directly relate to the environments that participants navigated in. Reporting these separately also removes ambiguity about what causes differences in symmetric modulation. When a single beta value is used to express modulation in the case of GLM analysis, it is influenced both by phase alignment and peak magnitudes independently, but typically interpreted as being modulated by phase alignment alone.

### Synthesized data

#### Simulation

The goal of the simulation was to generate a population of MEC neurons firing as a rodent moves around an open-field arena in order to match, and extrapolate beyond the limits of our empirical dataset. We focus on grid cells but also simulate head direction cells, conjunctive grid-x-head direction cells, spatially informative cells and pure noise cells. The ultimate goal being to do a directional analysis of the population signal with ground truth knowledge of individual neuron activity.

We use a stochastic model in which probabilities are sampled to give a spike count over a time interval. Cell populations are any combination of *grid, conjunctive* (grid x head direction, hd), *hd, place*, and *random* firing cells. While place cells are not typically found in the MEC, other spatially tuned/spatially informative cells are^2,24^. Although we call them “place cells” when considered collectively, in the model they are a proxy for the type of spatially dependent noise added to the grid cell signal from these spatially informative neurons of the MEC. In the model *place* and *pure grid* cells have only a spatial firing probability *p*(*x, y*), *hd* cells have only a direction firing probability *p*(θ), *conjunctive* cells have a joint probability *p*(*x, y*) *p*(θ), and *random* cells have a uniformly random probability of firing.

Firing is based on a stochastically determined firing rate, which is translated into a Bernoulli spike/no-spike probability of *n* trials where *n* is the firing rate multiplied by time interval. For this study we simulated sampling at 2 Hz (somewhere in the range of a very high fidelity BOLD signal), so the time interval was 0.5 s. A n-trial Bernoulli distribution is approximated by a Poisson distribution, which to speed up computation we further approximated with a gaussian distribution.

*Grid* and *place* cells were modeled as 2D normal distributions firing fields based on distance to the location of the field center. For *hd and conjunctive* cells directional firing probability was modeled as circular normal distribution based on circular distance to the tuning angle. If desired, grid patterns were generated such that they could have a random degree of ellipticity as well as a uniform amount of location noise with respect to the center of the field relative to the perfectly geometric pattern. Directional tuning for conjunctive cells was tuned to match the same tuning specificity (mvl) from the empirical distribution of the Gardner et al. 2022^55^ dataset (containing only grid and conjunctive grid-x-hd cells) at uniformly random tuning angles. This dataset is a subset of the Vollan et al. 2025^23^ dataset which had not been released at the time the model was being built. HD cells had uniform random mvl value from [0.7,1] with a uniformly random tuning angle.

Trajectories were generated as a collection of individual straight segments of fixed length and varying direction. Directions were sampled every n degrees with some added noise to avoid artifacts from sampling an exactly symmetric set of directions. At every direction the starting points of the segments were randomly distributed in an approximately circular shape forming disk shaped sample space collectively.

The overall parameters of the simulations usually matched a typical rodent experiment’s, including grid-spacing in the tens of centimeters, and trajectory area of around 1-2 *m*^2^. Path segment length was usually in the 10’s to 100’s of centimeters, with speed held constant at around 20 cm/s, translating to sampling at approximately every 10 cm. It should be noted that straight paths of this length are not typical in empirical rodent openfield experiments, and consequently are the main motivation for building this model as they are required to see the hexadirectional signal.

A typical simulation might have segment lengths of 150 cm × 50 segment samples per angle x 60 angles samples so ∼ 45,000 points sampled per cell. To optimize this we converted as much of these computations as we could to matrix multiplications and performed them on a single GPU with 14GB of VRAM. With this approach we could simulate 1 million cells for a total rodent distance of 4.5 km (∼6hr simulated time), in around 20 minutes, with the majority of simulation time determined by the number of cells. We validated the model by generating directional and spatial firing rate maps and checking the statistical distributions of our simulated data to see that we were generating spike trains and rate maps that would seem reasonable in a rodent experiment.

#### Hybrid approach

Despite developing a fairly detailed model with lots of parameters to explore, when the Vollan et al. 2025^23^ dataset became available, we decided that we wanted to draw our conclusions as much from this dataset as possible since it contains a wealth of simultaneously recorded cells in addition to grid cells, whose complex spatial and directional characteristics could not be replicated with simple stochastic exploration of our computational model’s parameter space. However despite having this excellent data to explore, it was imperative to have many long straight trajectory samples in all directions, which as mentioned, the rodents themselves did not provide. For this reason, we essentially lifted our trajectories from the simulation, and instead sampled from the empirically calculated directional *f*(θ_*t*_) and spatial *f*(*x* _*t*_, *y*_*t*_) firing maps, which we treated as independent firing-rate profiles whose product defines a separable joint firing-rate model. To correct for biases from the rodents sampling only a subset of the directional and 2D space in the empirical trajectories, we also normalize such that mean firing of the sampled trajectory matched the empirical mean firing rate 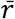 :

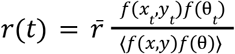

The assumption that spatial and directional tuning is independent is probably not entirely accurate. For example Gerlei et al. 2020^56^ reports that in pure grid direction cells, directional tuning changes per field. We did try more complex models of spatial and directional tuning dynamics to model individual cells, including GLMs and Gaussians processes fitted to the empirical *x, y*, θ, *r*(*t*), but ultimately we lacked enough data to cover this joint probability space dependently unless we used large spatial and directional bins, which did not yield enough specificity to generate realistic grid cell firing profiles.

Ultimately we decided that using this framework of factorized independent spatial and directional tuning was fairly in line with how these tunings are usually discussed in the field, and more relevantly, how they are utilized to construct the hypotheses that lead to the hexadirectional signal.

In this regime the ratemaps generated from our sampled trajectories match the original rate maps almost exactly, so conceptually we can say that we simply extend these first order dynamics for longer trajectories. That said, we are likely missing some relevant spatial and directional dependence or other higher-order temporal or network dynamics, but we are not aware of any hypotheses related to the hexadirectional signal that rely on these. Notably because we relied on the empirical spatial ratemaps, the length of our segments were still bounded to the confines of the original environment, a ∼1.5m square (with the exception of two recordings in a circular environment), and the empirical grid patterns (see Figure 4A middle, x-axis). Lastly we initially treated r(t) as a stochastic value λ in a poisson distribution, but found that with the number of samples we are taking per trajectory, 10s of thousands, this had negligible effect while slowing down the simulation noticeably. Instead stochasticity is achieved by randomly sampling trajectory segments.

### Modified grid score

While not central to this study, we made an attempt to better classify grid cells in our ∼24k cell database by modifying how we quantify gridness in spatial firing ratemaps. The canonical grid score method^52^ extracts a ring out of the rate map autocorrelogram, ideally at the radius and width to match a ring of autocorrelated grid fields. This ring is rotated and correlated/anticorrelated in alternating 30 degree increments. The most important modification to this method we made is to quantify the roundness of these firing fields in the autocorrelated image, and to more precisely identify them to extract a more accurate ring. We made this change because of the many false positive instances of high grid scores from noise ratemaps we had using the canonical method, the reasons of which are described below. We do this mostly with a rudimentary image processing pipeline.

#### Methodological reasons for false positives in gridness score

At the conceptual level, the false positives in the standard ‘gridness’ measure come from over-rewarding radial symmetry (6-fold in this case) which has a generally high chance probability in autocorrelograms, while under rewarding translational symmetry which is a defining characteristic of lattice patterns (the autocorrelation does accomplish this in theory, but it is ultimately washed out in downstream steps as described below). More specifically, these false positives occur from three factors. 1. The relatively small amount of the autocorrelogram that is correlated (area of the ring). 2. The relatively anisotropic pattern (random spatial peaks, not uniform noise) created by the pre-smoothing of the ratemaps and/or the inherent smoothing of the autocorrelation operation. 3. The fundamental mismatch between the geometry of the shape of the spatial bin being correlated (portion of ring when divided by 6, i.e. rounded trapezoid) and the shape of ideal grid fields themselves (ideally circular). At face value, the 30° increment correlation/anticorrelation procedure produces a maximal effect (a combined correlation value of 2) when a 60° degree slice of a ring shaped pattern inverts every 60° degrees. Consequently the part of the grid field that does not fit in the trapezoidal bin systematically detracts from the correlation magnitude. In our simulation of idealized perfect grid cells, this led to maximal grid scores of around 1.5. On the other hand, random ratemaps can often lead to inadvertent 6-fold symmetries, especially given their anisotropic nature, which would regularly lead to grid scores higher than this threshold.

#### A partial solution with a focus on firing field circularity using image processing

Given the mismatch of the geometry described above, we found that one requirement that was really impactful was identifying circular firing fields in the autocorrelated ratemaps. This eliminated a lot of false positives that arose from the relatively high levels of 6-fold spatial symmetry from autocorrelated, smoothed random firing. We did this by requiring some prior on expected grid field size in the form of a default grid field radius. We start by removing high frequency image noise (salt and pepper etc), and then use a round spatial filter based on this default field radius to enhance larger firing patterns, before autocorrelating the rate map. After this we apply a series of filters to discretize and threshold these larger firing patterns eventually leading to a binary map of large firing and no-firing regions. We next use some morphological operations (binary image processing methods) to clean up the image for small/jagged features. Ultimately we are left with a series of blobs. We then only consider fairly round blobs using the standard area/perimeter circularity measure, 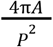. We then use the outer and inner radius of these blobs to form our ring, and apply the rest of the standard grid score measure on this binary image (30° increment correlation/anti-correlation). This addition was quite effective in analyzing this dataset, but in reality cannot be described as a universal method. That said, for those wishing to improve grid score, we highly recommend exploring this line of processing.

## Notes

### Competing Interest Statement

The authors have declared no competing interest.

### Summary of Updates

Mostly discussion. Some abstract, intro, and the title as well. Also another supplementary figure.

