## Supplementary figures for "Grid cells are not a parsimonious source of the hexadirectional signal"

### Figure S1. Pitfalls in hexadirectional analysis

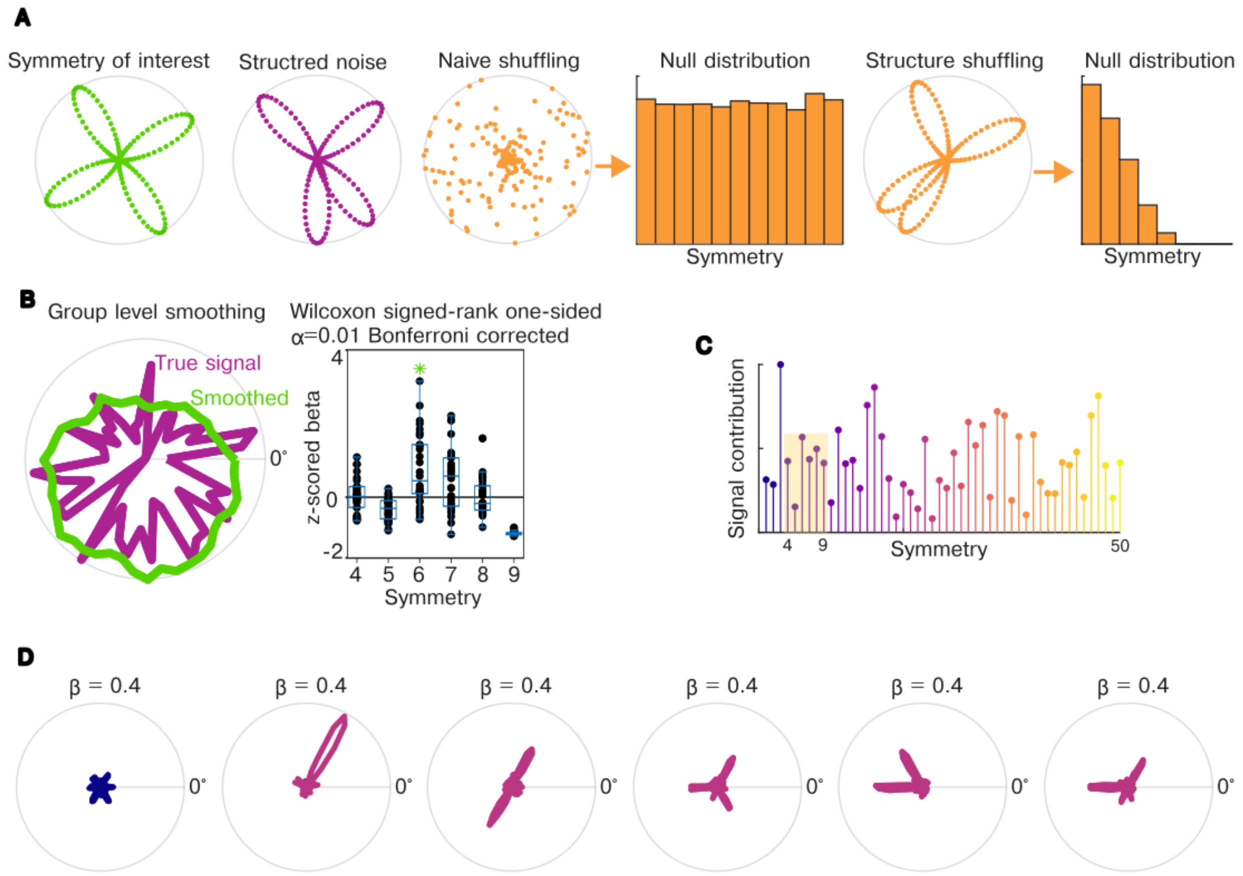


A. The importance of shuffling the correct feature of data. In this example we are testing for a 4-fold symmetry in data composed of 4 random directional peaks. In the first case we assume a null hypothesis where bin values are independent of each other. This naive shuffling of bin labels produces a uniform null distribution of symmetries. In the second case we assume an underlying structure to the data where nearby direction bins are correlated and thus shuffle peaks spanning multiple bins. This results in a much different null distribution of symmetries.

B. An illustrative simulation showing that optimizing parameters in the data processing can create a statistically significant results from noise. Here we simulate the data for 30 subjects sampled in 60 direction bins with uniform random numbers. We use a moving average to smooth the data, however we select window size to maximize 6 symmetric modulation at the group level. We then test statistical significance of the hexadirectional signal. This process achieves statistically significant hexadirectional modulation from random data frequently (every 1-3 simulations). In theory a moving average has zero gain at any harmonic k and window width w for which $k{\cdot}w/n$ is an integer, so a suitably chosen window can silence k = 4 and 8 outright while passing k = 6.

C. An illustrative plot showing that hexadirectional analysis typically only controls for a very small subset of possible directional symmetries [eg, 4-9], instead of the full spectrum (as limited by directional sampling precision). This is of concern because the lower number of control symmetries, the higher the chance of false positives. This plot shows the directional symmetry modulation for a random directional signal of 100 direction bins, note the variance in signal contribution amplitudes.

D. Misleading β's. An illustrative example of a pitfall related to quantifying hexadirectional signal with unbounded regression coefficients. In this example each signal has the same total energy (total firing), note r-axis magnitude. While the first case is truly hexadirectional, the rest are not, yet all produce the same regression coefficient.

### Figure S2. Effect of nonlinearity exponent and MEC population size on hexadirectional modulation

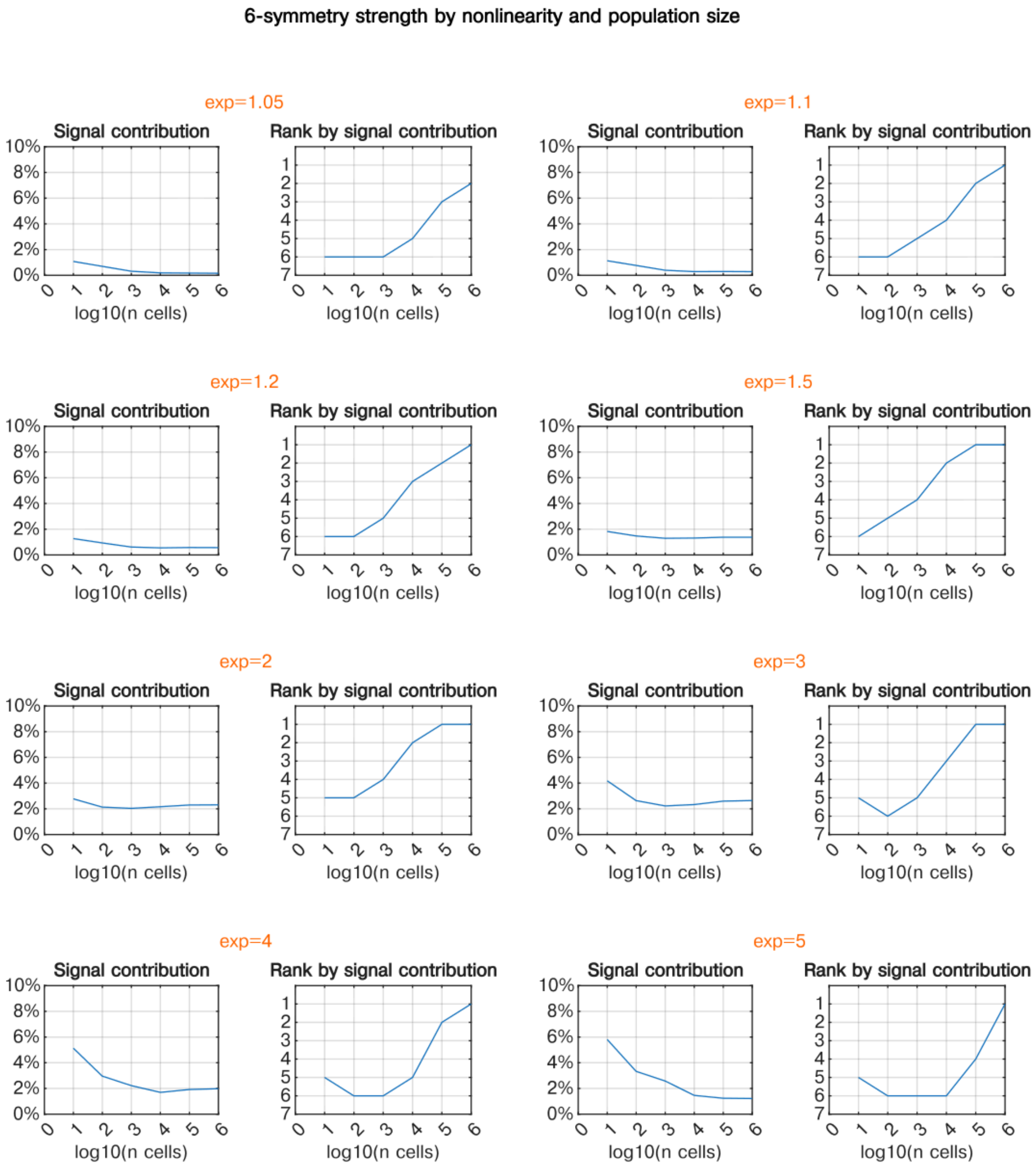


Here we explore two parameters, nonlinearity strength (power-function exponent), and population size on hexadirectional modulation using Monte Carlo simulation (100 simulations per population $n$). The same as Figure 3G but for additional exponents (see corresponding legend). Notably in this plot x-axis is expressed by the size of the total population, of which grid cells are roughly 15%. We also extend the simulation up to 1 million cell populations. Interestingly we see signal contribution at large populations first increase and then decrease as nonlinearity power increases (note: this is a subtle effect). The same appears true for rank (only intermediate nonlinearities each rank 1 at population size $n = 10^5$ cells). Decrease in signal in the larger nonlinearities is likely due to decrease in anisotropic directional noise as directional noise becomes more uniformly distributed, removing energy from other symmetries, simultaneously as the 0 symmetry (mean, which is not removed from the signal during analysis, see Methods), the majority of the signal, increases nonlinearly with population size. This is supported by the fact that though the signal is high at low population sizes, the rank is not, indicating that all symmetries are higher relative to the mean.

### Figure S3. Post-hoc analysis flexibility increases the probability of apparently selective sixfold effects under a pure null

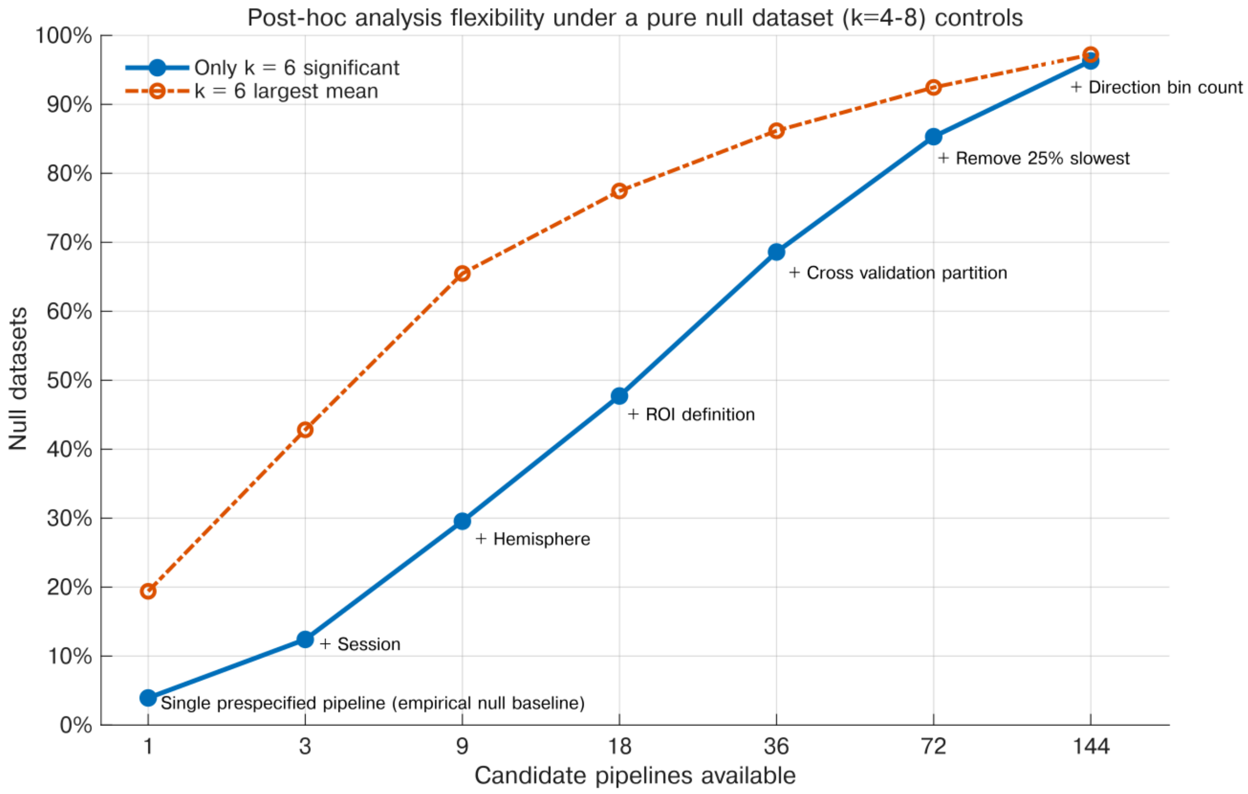


We simulated 5,000 cohorts of 20 participants, with 500 samples per participant, random movement directions, and Gaussian activity containing no directional or sixfold structure. Each fixed dataset was re-analysed while progressively adding plausible researcher degrees of freedom: session (all data, first session, second session), hemisphere (bilateral, left, right), ROI definition (two overlapping synthetic ROIs), cross-validation partition (first-half/second-half, odd/even samples), speed-based exclusion (all samples, remove 25% slowest; modeled under the null as a random 25% subset independent of activity), and direction-bin count (24, 30 bins). For each available parameter space, the pipeline where only k=6 significant was selected. Curves show the proportion of null cohorts in which only k=6 was significant while (k=4,5,7,8) were not, and in which k=6 had the largest cohort mean among (k=4–8). For each pipeline, we used the canonical train/test hexadirectional procedure: the preferred k-fold phase was estimated in training data and then held fixed while modulation amplitude was evaluated in independent held-out data ($\beta_k$). Group-level significance was assessed by testing whether the held-out modulation coefficients ($\beta_k$) were greater than zero across participants using a one-sided one-sample t-test ($p<0.05$ $p<0.05$).
